# The Requirement For US28 During Cytomegalovirus Latency Is Independent Of US27 And US29 Gene Expression

**DOI:** 10.1101/2020.01.19.911503

**Authors:** Benjamin A. Krishna, Amanda B. Wass, Rajashri Sridharan, Christine M. O’Connor

## Abstract

The ability to establish a latent infection with periodic reactivation events ensures herpesviruses, like human cytomegalovirus (HCMV), lifelong infection and serial passage. The host-pathogen relationship throughout HCMV latency is complex, though both cellular and viral factors influence the equilibrium between latent and lytic infection. We and others have shown one of the viral-encoded G protein-coupled receptors, US28, is required for HCMV latency. US28 potentiates signals both constitutively and in response to ligand binding, and we previously showed deletion of the ligand binding domain or mutation of the G protein-coupling domain results in the failure to maintain latency similar to deletion of the entire US28 open reading frame (ORF). Interestingly, a recent publication detailed an altered phenotype from that previously reported, showing US28 is required for viral reactivation rather than latency, suggesting the US28 ORF deletion impacts transcription of the surrounding genes. Here, we show an independently generated US28-stop mutant, like the US28 ORF deletion mutant, fails to maintain latency in hematopoietic cells. Further, we found *US27* and *US29* transcription in each of these mutants was comparable to their expression during wild type infection, suggesting neither US28 mutant alters mRNA levels of the surrounding genes. Finally, infection with a US28 ORF deletion virus expressed US27 protein comparable to its expression following wild type infection. In sum, our new data strongly support previous findings from our lab and others, detailing a requirement for US28 during HCMV latent infection.

## 1 Introduction

Human cytomegalovirus (HCMV) is a ubiquitous pathogen that latently infects the majority of the population (Khanna and Diamond, 2006). Latent infection in healthy individuals rarely poses a significant health risk, however immune dysregulation can lead to reactivation and CMV-associated disease, which can be fatal (Arvin et al., 2004; Ramanan and Razonable, 2013; Griffiths et al., 2015; Ljungman et al., 2017). This underscores the need to better understand these phases of viral infection to prevent disease and improve patient outcomes.

Our current understanding of the biological mechanisms controlling latency and reactivation remain incomplete, though work from many labs have detailed the importance of both host and viral factors in these processes (Collins-McMillen et al., 2018; Elder and Sinclair, 2019). We and others have shown the viral G protein-coupled receptor (GPCR) US28 is required for viral latency (Humby and O’Connor, 2015; Wu and Miller, 2016; Krishna et al., 2017; Krishna et al., 2019). US28 is expressed during both latent and lytic infection (Krishna et al., 2018), and we were the first to detail its requirement for successful HCMV latent infection (Humby and O’Connor, 2015). Building upon our original work, we more recently showed US28 regulates the expression and activity of cellular fos (c-fos) (Krishna et al., 2019), a component of the activator protein-1 (AP-1) transcription factor complex (Halazonetis et al., 1988). US28’s attenuation of c-fos leads to a decrease in AP-1 binding to the major immediate early promoter (MIEP) (Krishna et al., 2019), a key regulator in the latent-to-lytic switch (Collins-McMillen et al., 2018). Additionally, our data revealed a requirement for G protein-coupling, and to a lesser extent, ligand binding to US28, suggesting US28-mediated signaling is important for this phenotype (Krishna et al., 2019). This is consistent with findings from the Sinclair Lab, who showed US28 is required for latency in monocytes. Their work also detailed specific signaling pathways US28 impacts to ensure MIEP silencing in these latently-infected cells (Krishna et al., 2017; Elder et al., 2019). Finally, Wu and Miller showed infection of THP-1 monocytes with a US28-deletion mutant resulted in robust IE1/2 protein expression, compared to cultures infected with virus expressing US28 (Wu and Miller, 2016). In sum, these data strongly support a significant role for US28-mediated signaling in maintaining a latent infection in hematopoietic cells.

Recent work from Crawford et al. challenges these previous findings, as they show US28 is not required for latency, but rather is necessary for reactivation. Using a US28 stop mutant, the investigators demonstrated latency was maintained, but the infection failed to reactivate following the addition of stimuli. Surprisingly, they showed infection with mutant virus containing a point mutation in the ligand binding domain of US28 (Y16F) failed to maintain latent infection, suggesting that while US28 protein (pUS28) expression is not required, ligand binding is essential for latent infection. Crawford et al. suggested a compelling argument that the differences between their work and others were due to the complete ORF deletion of US28 other groups had performed, positing the US28 ORF deletion could impact wild type expression of surrounding genes, namely *US27* and *US29* (Crawford et al., 2019). To date, the impact of the complete US28 ORF on *US27* and *US29* remains unknown.

As this is a legitimate concern, we generated an additional set of viral recombinants using an additional BAC-derived clinical isolate, FIX (BFX*wt*-GFP; *wt*). We constructed a triple flag-tagged US28 recombinant (BFX-GFP*in*US28-3xF; *in*US28-3xF), from which we then inserted a stop codon immediately following the first methionine (BFX-GFP*stop*US28; *stop*US28). Using these independently-generated viruses, we now show, consistent with our previous work (Humby and O’Connor, 2015; Krishna et al., 2019), US28 is dispensable for efficient lytic viral growth in fibroblasts, however this viral GPCR is essential for latency. Our data herein aligns with previous work (Humby and O’Connor, 2015; Wu and Miller, 2016; Krishna et al., 2017; Krishna et al., 2019), as ablation of US28 protein expression in hematopoietic cells that support latency results instead in a lytic-like infection. Additionally, we assessed *US27* and *US29* transcription in both *stop*US28- and TB40/E*mCherry*-US28Δ (US28Δ)-infected fibroblasts, as well as US27 protein (pUS27) in cells infected with a US28Δ variant and found deletion of the entire open reading frame (ORF) did not impact these transcripts or translation of pUS27 in the absence of US28 expression. Together, our data confirm the requirement for US28 in maintaining HCMV latency, which is independent of *US27* or *US29* mRNA expression.

## 2 Materials and Methods

### 2.1 Cells & Viruses

Primary human foreskin fibroblasts (HFF, passages 9 to 13), MRC-5 embryonic lung fibroblasts (MRC-5, passages 21 to 30; ATCC, cat#CCL-171, RRID: CVCL_0440), or newborn human foreskin fibroblasts (NuFF-1, passages 13 to 25; GlobalStem, cat#GSC3002) were maintained in Dulbecco’s modified Eagle medium (DMEM), supplemented with 10% fetal bovine serum (FBS), 2 mM L-glutamine, 0.1 mM nonessential amino acids, 10 mM HEPES, and 100 U/ml each of penicillin and streptomycin. Kasumi-3 cells (ATCC CRL-2725, RRID: CVCL_0612) were maintained in RPMI 1640 medium (ATCC, cat#30-2001), supplemented with 20% FBS, 100 U/ml each of penicillin and streptomycin, and 100 μg/ml gentamicin at a density of 5 × 10^5^ to 1 × 10^6^ cells/ml. Murine stromal cells S1/S1 and M2-10B4 (MG3) were kind gifts from Terry Fox Laboratories, BC Cancer Agency (Vancouver, BC, Canada). S1/S1 cells were maintained in Iscove’s modified Dulbecco’s medium (IMDM), supplemented with 10% FBS, 1 mM sodium pyruvate, and 100 U/ml each of penicillin and streptomycin. MG3 cells were maintained in RPMI 1640, supplemented with 10% FBS and 100 U/ml each of penicillin and streptomycin. S1/S1 and MG3 cells were plated in a 1:1 ratio (~1.5 × 10^5^ cells of each cell type) onto collagen-coated (1 mg/ml) 6-well plates in human CD34^+^ long-term culture media (hLTCM), containing MyeloCult H5100 (Stem Cell Technologies, cat#5150) supplemented with 1 μM hydrocortisone, and 100 U/ml each of penicillin and streptomycin. The next day, the cells were irradiated using a fixed source ^137^Cesium, Shepherd Mark I Irradiator at 20 Gy, after which the cells were washed three times with 1X PBS, then resuspended in fresh hLTCM and returned to culture. Irradiated murine stromal cells were utilized the following day as feeder cells for the primary CD34^+^ hematopoietic progenitor cells (HPCs). Primary CD34^+^ HPCs were isolated from de-identified cord blood samples (Abraham J. & Phyllis Katz Cord Blood Foundation *d.b.a.* Cleveland Cord Blood Center & Volunteer Donating Communities in Cleveland and Atlanta) by magnetic separation, as described elsewhere (Umashankar and Goodrum, 2014). Isolation and culture methods for the primary CD34^+^ HPCs are detailed below. All cells were maintained at 37°C/5% CO_2_.

HCMV bacterial artificial chromosome (BAC)-derived strain TB40/E (clone 4) (Sinzger et al., 2008) previously engineered to express mCherry to monitor infection, TB40/E*mCherry* (O’Connor and Shenk, 2011), was used in this study. TB40/E*mCherry*-US28-3xF, TB40/E*mCherry*-US28Δ were previously characterized (Miller et al., 2012). An additional BAC-derived isolate engineered to express eGFP as a marker of infection, BFX*wt*-GFP (Murphy et al., 2008), was used to generate a recombinant virus expressing a US28 C-terminal triple FLAG epitope tag, BFX*wt*-GFP-US28-3xF, by bacterial recombineering techniques described in elsewhere (O’Connor and Miller, 2014). Briefly, the 3xF epitope and Kan-frt cassette were PCR amplified from pGTE-3xFLAG-Kan-frt (O’Connor and Shenk, 2011) using the 3xF-Kan-frt insertion primers (Supplementary Table 1). This product was then used to generate BFX-GFP-*in*US28-3xF by recombination (e.g. ref. (O’Connor and Shenk, 2011). BFX*wt*-GFP-*in*US28-3xF was then used to generate two independent US28 stop mutants using *galK* recombineering, as described previously (O’Connor and Miller, 2014). Briefly, the *galK* gene was amplified by PCR using primers listed in Supplementary Table 1. Recombination-competent SW105 *Escherichia coli* containing BFX-GFP-*in*US28-3xF were transformed with the resulting PCR product. *GalK*-positive clones were selected and electroporated with the double stranded reversion oligoucleotide (Supplementary Table 1) and mutants were counter-selected against *galK*. Two independently generated mutants, BFX-GFP-*stop*US28-S1 and BFX-GFP-*stop*US28-S2, were validated by Sanger sequencing. The multiple epitope tag viral GPCR mutant was generated using TB40/E*mCherry*-US28-3xF as a backbone. Each of the remaining three viral GPCRs were serially epitope tagged with the primers in Supplementary Table 1, and recombinant clones were sequenced following each reversion. The resulting virus, multi-tag vGPCR (vGPCR*multi*), contains the following epitope tags: US28-3xF, US27-3xHA, UL33-c-myc, and UL78-V5. vGPCR*multi* was then used to generate vGPCR*multi*-US28Δ using *galK* recombineering techniques. The primers used to generate this mutant are previously described (Miller et al., 2012). The sequence for vGPCR*multi*-US28Δ was verified by Sanger sequencing. All viral stocks were propagated and titered by 50% tissue culture infectious dose (TCID_50_) as described (e.g. ref. (O’Connor and Shenk, 2012).

### 2.2 Viral Growth Analyses

Multi-step growth assays were performed using fibroblasts (MRC-5, NuFF-1) by infecting cells at a multiplicity of infection (moi) of 0.01 TCID_50_/cell. Infectious supernatants were collected over a time course of infection and stored at −80°C until processing. Infectious virus was then titrated on naïve fibroblasts (MRC-5, NuFF-1) and analyzed by TCID_50_ assay.

### 2.3 Viral RNA & Protein Assays

For viral transcript analyses, primary NuFF-1 fibroblasts were infected at an moi = 0.5 TCID_50_/cell. Total RNA was collected 96 hours post-infection (hpi) and RNA was extracted with the High Pure RNA Isolation kit (Roche, cat#11828665001), according to the manufacturer’s instructions. cDNA was generated from 1.0 μg of RNA using TaqMan Reverse Transcription (RT) Reagents and random hexamer primers (Roche, cat#N8080234). Equal volumes of cDNA were used for quantitative PCR (qPCR) using gene specific primers, and cellular *GAPDH* was used as a control (Supplementary Table 1). Transcript abundance was calculated using a standard curve using 10-fold serial dilutions of a BAC-standard that also contains GAPDH sequence. Viral gene abundance was normalized to *GAPDH* for each sample. Each primer set had a similar linear range of detection for the BAC-standard (linear between 10^9^ and 10^4^ copies; *r^2^* > 0.95 for all experiments). Samples were analyzed in triplicate using a 96-well format CFX Connect (BioRad).

For immunofluorescence assays (IFA), primary MRC-5 or NuFF-1 fibroblasts were grown on coverslips and infected (moi = 0.5) as indicated in the text. Cells were harvested and processed as described elsewhere (e.g. refs. (O’Connor and Shenk, 2011; O’Connor and Murphy, 2012). Antibodies used include: anti-FLAG M2 (Sigma, cat#F3165, RRID:AB_259529; 1:1,000), anti-HA (Roche, cat#11867423001, RRID:AB_390918; 1:1,000), anti-c-Myc (Sigma, cat#M4439, RRID:AB_439694; 1:500), anti-V5 (Sigma, cat#V8137, RRID:AB_261889; 1;1,000), Alexa 488-conjugated anti-rat (Abcam, cat#ab150157, RRID:AB_2722511; 1:1,000), Alexa 488-conjugated anti-mouse (Fisher, cat#A11001, RRID:AB_2534069; 1:1,000), Alexa 488-conjugated anti-rabbit (Fisher, cat#A11008, RRID:AB_143165; 1:1,000), Alexa 647-conjugated anti-mouse (Abcam, cat#ab150115, RRID:AB_2687948; 1:1,000), 4’-6’-diamidino-2-phenylindole (DAPI). Coverslips were mounted onto slides with Slow-Fade reagent (Invitrogen, cat#S2828) or FluorSave Reagent (Calbiochem, cat#345789), and images were collected using a Zeiss LSM 510 or Leica SP8 confocal microscope.

To assess protein expression by immunoblot, ~3.0 × 10^5^ NuFF-1 fibroblasts were infected (moi = 0.5) for 96h. Cells were harvested in RIPA buffer, and equal amounts of protein were analyzed using the following antibodies: anti-FLAG M2 (Sigma, cat#F3165, RRID:AB_259529; 1:7,500), anti-IE1 (clone 1B12 (Zhu et al., 1995); 1:100), anti-pp65 (clone 8A8 (Bechtel and Shenk, 2002); 1:100), anti-HA (Roche, cat#11867423001, RRID:AB_390918; 1:1,000), anti-actin (Sigma, cat#A3854, RRID:AB_262011; 1:20,000), and goat-anti-mouse (cat#115-035-003, RRID:AB_10015289) or goat-anti-rat (cat#112-035-003, RRID:AB_2338128) horseradish peroxidase (HRP) secondary (Jackson ImmunoResearch Labs; 1:10,000).

### 2.4 Latency Infection & Extreme Limiting Dilution Assay

Kasumi-3 cells (moi = 1.0) were infected as described previously (e.g. ref. (Krishna et al., 2019). Briefly, cells were cultured in serum-low media (XVIVO-15; Lonza, cat#04-418Q) for 48h prior to infection. Kasumi-3 cells were infected at a density of 5.0 × 10^5^ cells/ml by centrifugal enhancement. At 7 days post-infection (dpi), cultures were treated with 20nM 12-O-tetredecanoylphorbol-13-acetate (TPA) or vehicle (DMSO) for an additional 2d. Infectious particle production was assessed by Extreme Limiting Dilution Assay (ELDA) on naïve NuFF-1 fibroblasts, as described previously (Umashankar and Goodrum, 2014).

Primary CD34^+^ HPC culture and infection conditions are described elsewhere (Umashankar and Goodrum, 2014). Briefly, CD34^+^ HPCs (moi = 2.0) were infected by centrifugal enhancement, followed by overnight incubation. Cells were washed and cultured over irradiated MG3:S1/S1 murine stromal cells (plated at 1:1 ratio, see above). At 7 dpi, a portion of each infected cell population was cultured in reactivation media (RPMI 1640, containing 20% FBS, 10 mM HEPES, 1 mM sodium pyruvate, 2 mM L-glutamine, 0.1 mM nonessential amino acids, 100 U/ml each penicillin and streptomycin, with 15 ng/ml each (all from R&D Systems): IL-6, G-CSF, GM-CSF, IL-3) or maintained in hLTCM. Infectious particle production was assessed by ELDA on naïve NuFF-1 fibroblasts, as detailed elsewhere (Umashankar and Goodrum, 2014).

## 3 Results

### 3.1 *stop*US28 virus replicates to wild type titers in fibroblasts

To ensure US28’s function is due to the absence of only this protein as opposed to potential off-site consequences resulting from deletion of the US28 ORF, we generated a new panel of recombinants using the BAC-derived, clinical isolate, BFX*wt*-GFP (*wt*) (Murphy et al., 2008). The first variant, BFX-GFP*in*US28-3xF (*in*US28-3xF) expresses a pUS28 fusion protein with three, tandem FLAG epitope repeats in the C-terminus of the protein (Fig 1A). Similar to our work in another BAC-derived clinical isolate, TB40/E, in which we made an identical tagged pUS28 recombinant virus (Miller et al., 2012), we observed robust pUS28 expression following lytic infection of fibroblasts by both immunofluorescence assay (IFA; Fig 1B) and immunoblot (Fig 1C). In line with previously published data (Slinger et al., 2010; Noriega et al., 2014), pUS28 localizes to a perinuclear region of infected fibroblasts, consistent with the assembly complex (Silva et al., 2003). Next, we generated two independently derived BFX-GFP*stop*US28 constructs using the *in*US28-3xF backbone (Fig 1A), allowing us to confirm protein ablation by both western blot analysis (Fig 1B) and IFA (Fig 1C). Further, consistent with previously published work (Dunn et al., 2003; Yu et al., 2003; Miller et al., 2012; Humby and O’Connor, 2015), we found pUS28 is not required for efficient viral replication in lytically infected fibroblasts, as the two, independently generated stop mutants grew to wild type titers (Supplementary Fig 1). Together, these data confirm pUS28 expression is ablated in *stop*US28-infected fibroblasts, and pUS28 is dispensable for lytic replication in these cells.

**Figure 1.**
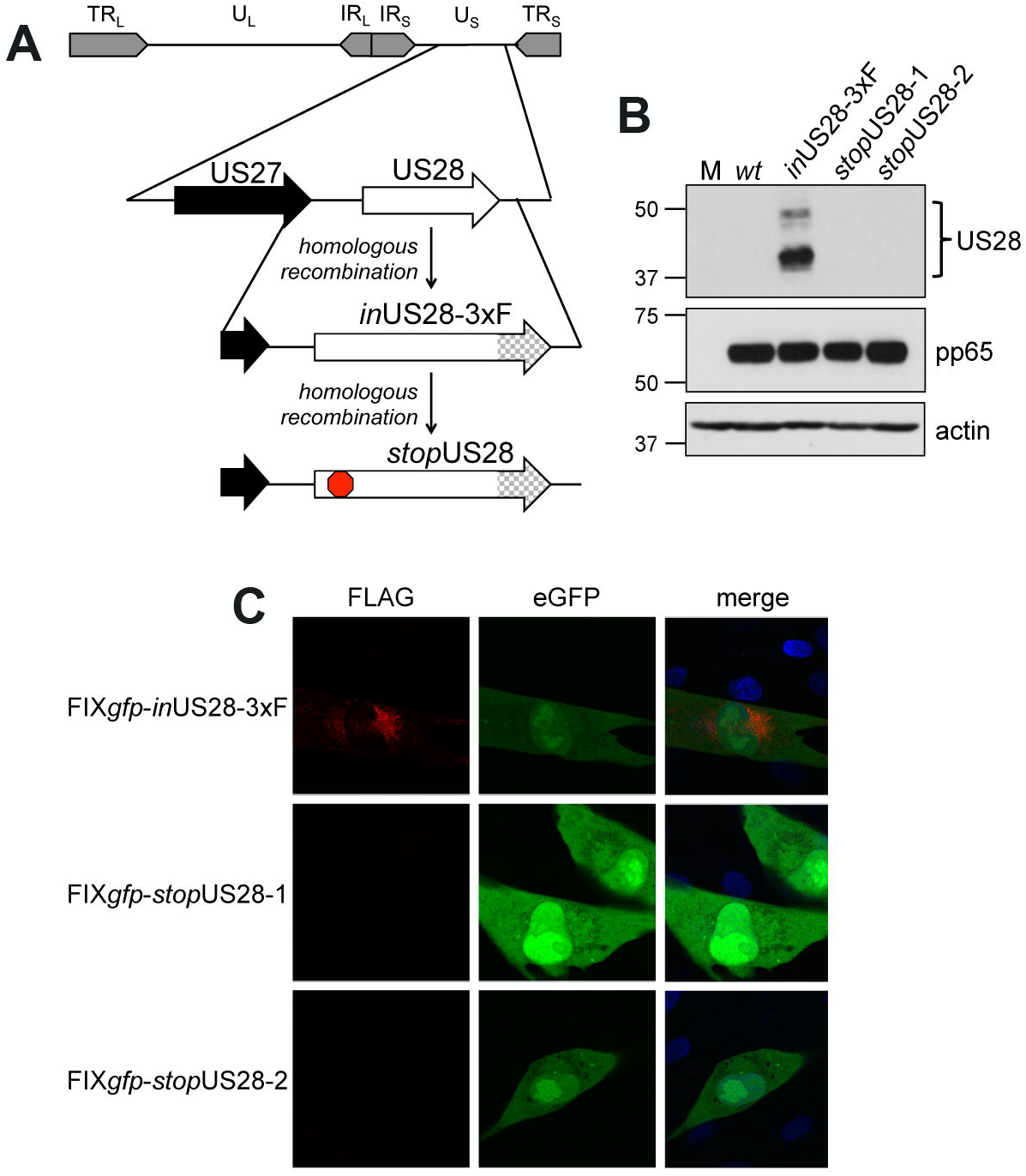
US28 protein expression is ablated in *stop* US28-infected fibroblasts. **(A)** BFX*wt*-GFP (*wt*) was used to generate *in*US28-3xF, which contains an in-frame triple FLAG epitope tag (3xF) at the C-terminal end of the ORF (checked arrow). *in*US28-3xF was then used as the template to generate two independent stop mutants, *stop*US28-1 and *stop*US28-2, that each contains a stop codon following the first methionine (red stop sign). **(B,C)** Fibroblasts were infected as indicated (moi = 0.5). **(B)** Cell lysates were harvested 96 hpi for immunoblot. α-FLAG was used to detect pUS28 expression, α-pp65 is a marker of infection, and actin is shown as a loading control. **(C)** Infected cultures were processed for IFA 72hpi. α-FLAG was used to detect pUS28 expression via the 3xF epitope (red). eGFP (green) is a marker of infection, and nuclei were visualized using DAPI (blue). Images were acquired using a 60x objective. **(B,C)** Representative images are shown; n = 3.

### 3.2 *stop*US28 fails to maintain latency in hematopoietic cells

We and others have shown deletion of the entire US28 ORF from TB40/E (Humby and O’Connor, 2015; Krishna et al., 2019), Titan (Krishna et al., 2017), and FIX (Wu and Miller, 2016) strains of HCMV results in the failure to establish/maintain latent infection of hematopoietic cells, including Kasumi-3 (Humby and O’Connor, 2015; Krishna et al., 2019) and THP-1 cell lines (Wu and Miller, 2016; Krishna et al., 2017; Krishna et al., 2019), as well as primary monocytes (Krishna et al., 2017) and cord blood-derived CD34^+^ HPCs (Krishna et al., 2019). However, recently published findings reported a US28 stop mutant in TB40/E is capable of maintaining latency in primary fetal liver-derived CD34^+^ HPCs, although this virus failed to reactivate (Crawford et al., 2019). The authors posited that complete ORF deletion possibly impacted efficient expression of surrounding genes, thus potentially contributing to the discrepancies in phenotypes between this and previous studies (Crawford et al., 2019). To determine if our newly generated US28 stop mutant displayed a similar phenotype during latent infection, we infected Kasumi-3 and cord blood-derived CD34^+^ cells with *wt* or *stop*US28 for 7d under latent conditions. We then divided each infected culture, treating half with reactivation stimuli for an additional 2d, where we treated Kasumi-3-infected cultures with TPA and cultured primary CD34^+^ HPCs in reactivation media. We then quantified the production of infectious particles by extreme limiting dilution assay (ELDA) on naïve fibroblasts. *stop*US28-infected cells failed to maintain a latent infection, as Kasumi-3 or CD34^+^ cells infected with this mutant produced infectious virus regardless of reactivation stimuli treatment (Fig 2). These data suggest ablating pUS28 by either introduction of a stop codon or deletion of the ORF results in a variant incapable of maintaining latency in hematopoietic cells.

**Figure 2.**
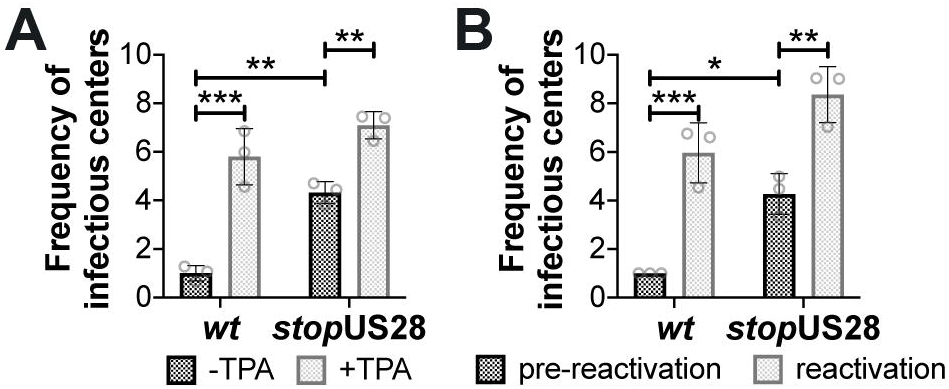
*stop*US28 fails to maintain latency in hematopoietic cells. **(A)** Kasumi-3 cells (moi = 1.0) or **(B)** CD34^+^ HPCs (moi = 2.0) were infected under latent conditions with the indicated viruses. At 7 dpi, infected **(A)** Kasumi-3 cells were treated with vehicle (DMSO; -TPA, black bars) or TPA (+TPA, gray bars), and **(B)** CD34^+^ HPCs were cultured in hLTCM (pre-reactivation, black bars) or reactivation media (reactivation, gray bars). The fold-change in the frequency of infectious particle production was quantified by ELDA on naïve fibroblasts 14 d later and is graphed relative to WT **(A)** -TPA or **(B)** pre-reactivation. Each data point (circles) is the mean of three technical replicates (i.e. one biological replicate). Error bars indicate standard deviation of three biological replicates. Statistical significance was calculated using two-way ANOVA analyses followed by Tukey’s post-hoc analyses. * *p* < 0.05, ** *p* < 0.01, *** *p* < 0.001

### 3.3 Deletion of the US28 ORF does not impact *US27* or *US29* transcription or US27 protein expression

While we observed no difference in the outcome of a US28 stop mutant versus a US28 ORF deletion mutant, we were concerned this mutation may affect neighboring viral transcripts, such as *US27*, which is encoded along with *US28* as a polycistronic transcript (Balazs et al., 2017). Thus, to ensure *US27* and *US29* mRNA expression are unaffected by altered pUS28 expression, we assessed each of these transcripts following lytic infection of fibroblasts with BFX*wt*-GFP or BFX*stop*US28, as well as TB40/E*mCherry* or TB40/E*mCherry*-US28Δ. We chose to evaluate these transcripts during the lytic life cycle because neither of these genes is expressed during latency (Humby and O’Connor, 2015; Cheng et al., 2017; Shnayder et al., 2018). To this end, we lytically infected fibroblasts (moi = 0.5), harvested total RNA at 96 hpi, and performed RTqPCR to quantify *US27* and *US29* transcripts, as well as *US28*, *UL123*, and *UL99* as controls. We found ablating pUS28 expression did not impact the transcription of *US27* or *US29* in either US28 recombinant virus (Fig 3). Since *US27* and *US28* originate from a polycistronic RNA, we also assessed US27 protein (pUS27) expression in the context of US28 ORF deletion. To this end, we generated a virus construct in the TB40/E*mCherry* background that contains a different epitope tag on the C-terminus of each viral-encoded GPCR, termed TB40/E*mCherry*-vGPCR*multi* (vGPCR*multi*). Each vGPCR is tagged as follows: US27-3xHA, US28-3xF, UL33-myc, and UL78-V5 (Supplementary Fig 2A). Using this construct, we then generated a US28 deletion, including the triple FLAG epitope tag, termed vGPCR*multi*-US28Δ (Supplementary Fig 2A), which replicated with wild type kinetics (Supplementary Fig 2B). We then used these newly-generated viral recombinants to lytically infect fibroblasts (moi = 0.5) to determine their localization and expression by IFA and immunoblot, respectively. vGPCR*multi*-infected fibroblasts express each of the four vGPCRs (Supplementary Fig 3), while vGPCR*multi*-US28Δ fails to express pUS28, as expected (Fig 4). Importantly, complete ORF deletion of US28 does not impact pUS27 expression (Fig 4A) or localization (Fig 4B), consistent with our transcriptional data (Fig 3). Together, these data suggest ablation of pUS28 expression does not impact *US27* and *US29* expression.

**Figure 3.**
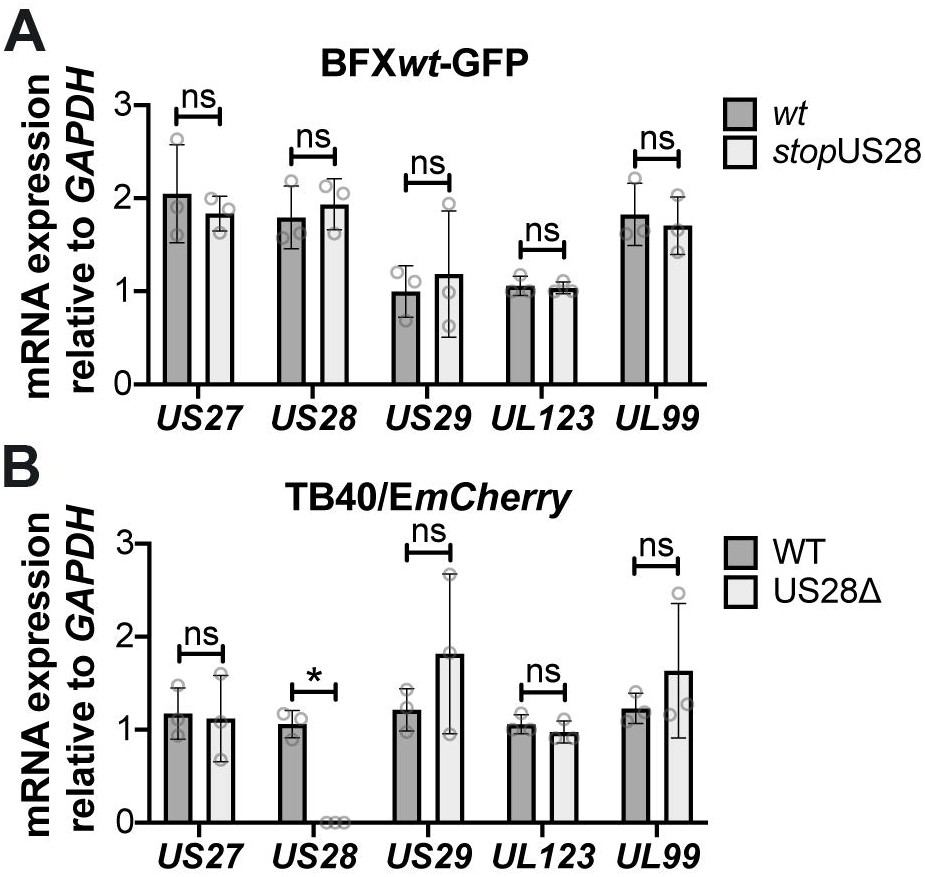
Abrogating pUS28 expression does not impact *US27* or *US29* transcription. NuFF-1 fibroblasts were infected (moi = 0.5) with **(A)** BFX*wt*-based or **(B)** TB40/E*mCherry*-based viruses. Total RNA was harvested 96 hpi and *US27*, *US28*, *US29*, *UL123*, and *UL99* mRNA levels were quantified by RTqPCR. Viral gene expression is plotted relative to cellular *GAPDH*. Each data point (circles) is the mean of three technical replicates (e.g. one biological replicate). Error bars indicate standard deviation of three biological replicates, and statistical significance was calculated using two-way ANOVA analyses followed by Tukey’s post-hoc analyses. * *p* < 0.05; ns, not significant.

**Figure 4.**
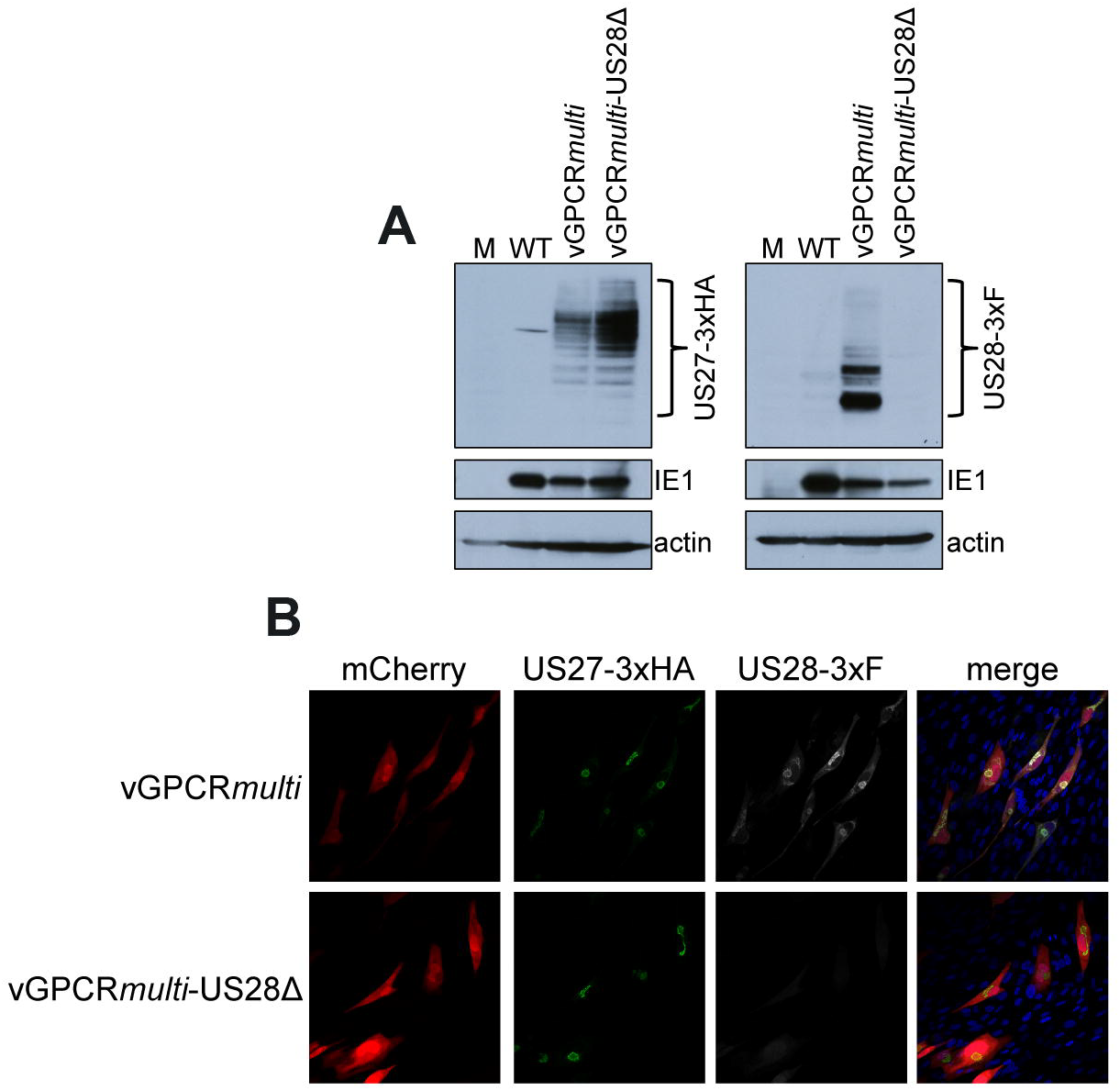
US28Δ-infected fibroblasts display wild type pUS27 levels. NuFF-1 fibroblasts were mock-infected (M) or infected with TB40/E*mCherry* (WT), TB40/E*mCherry*-vGPCR*multi*, or TB40/E*mCherry*-vGPCR*multi*-US28Δ (moi = 0.5). At 96 hpi, **(A)** cell lysates were collected for immunoblot or **(B)** harvested for IFA. **(A,B)** α-FLAG was used to detect US28 via the 3xF epitope, α-HA was used to detect US27 via the 3xHA epitope. **(A)** IE1 is shown as a marker of infection, and cellular actin serves as a loading control. **(B)** US27-3xHA (green), US28-3xF (white), mCherry (red) serves as a marker of infection. DAPI (blue) was used to visualize nuclei. Images were acquired using 40x objective. **(A,B)** Representative images are shown (n = 3).

## 4 Discussion

Our findings using newly generated recombinant HCMV constructs reveal pUS28 expression is required for HCMV latency. Our data confirm previous work, wherein we and other groups demonstrated US28 ORF deletion viruses favor lytic rather than latent infection in hematopoietic cells. We now show the insertion of a stop codon after the first methionine in the US28 ORF in the BFX*wt*-GFP background ablates protein expression, which, similar to the US28 ORF deletion virus we previously generated in the TB40/E background, results in a lytic-like infection of both Kasumi-3 and cord blood-derived CD34^+^ cells. Our data also suggest *US27* and *US29* gene expression are not impacted by the lack of pUS28 expression, as the US28 stop and deletion viruses express these neighboring transcripts to wild type levels during lytic infection. Furthermore, we show deleting the US28 ORF does not alter the localization or expression of pUS27. Together, our findings support previously published findings detailing the requirement of pUS28 expression for HCMV latency in hematopoietic lineage cells.

As mentioned, recent work from Crawford et al. showed the insertion of two tandem stop codons following the first methionine in the US28 ORF resulted in a TB40/E-based mutant capable of maintaining latency, yet incapable of reactivating in response to stimuli. While the US28 stop mutant maintained latent infection, a point mutation (Y16F) within the US28 ligand binding domain failed to do so, leading the authors to conclude that while pUS28 expression was dispensable, ligand binding to pUS28 was required for viral latency (Crawford et al., 2019). Interestingly, mutating the US28 G protein-coupling domain, or the canonical ‘DRY’ motif, which renders US28 “signaling dead” (Waldhoer et al., 2002; Maussang et al., 2006; Maussang et al., 2009; Miller et al., 2012), phenotypically resembled the US28 stop mutant (Crawford et al., 2019). Whether signaling constitutively or in response to ligand binding, a functional G protein-coupling domain is required to potentiate downstream signaling (Haskell et al., 1999; Auger et al., 2002; Schwartz et al., 2006; Rovati et al., 2007). Thus, perhaps in their system, US28 is not behaving as a canonical GPCR. We previously demonstrated a requirement for both US28’s G protein-coupling domain, and to a lesser extent the ligand binding domain, in the context of latent infection. This revealed US28-mediated signaling is required for viral latency and is at least partly dependent upon US28’s interaction with ligand(s) (Krishna et al., 2019). Work from the Sinclair Lab also detailed the requirement for US28’s G protein-coupling domain in suppressing IE protein expression in THP-1 cells, though the Y16F ligand binding mutant suppressed IE protein expression to wild type levels (Krishna et al., 2017). It is important to note we generated a ligand binding mutant, US28ΔN, in which we deleted amino acids 2-16 in the US28 ORF (Krishna et al., 2019), in contrast to the single point mutation, Y16F. We chose to delete these amino acids, as Casarosa et al. previously showed chemokines differentially bind to specific residues within pUS28’s N-terminus (Casarosa et al., 2005). Specific point mutants within this region of pUS28 will undoubtedly prove useful towards identifying the specific ligand(s) with which pUS28 interacts to potentiate latency-specific signaling. Nonetheless, this important variance in the mutants could explain some distinctions between the aforementioned work and ours.

What other differences might account for the distinct findings mentioned above? HCMV latency and reactivation are not trivial phases of infection to study in tissue culture. There are various culture systems, viral backgrounds, and culture conditions (e.g. media and additives) that could impact results. We showed pUS28 is required for latency using both the TB40/E (Humby and O’Connor, 2015; Krishna et al., 2019) and BFX*wt* backgrounds, using either ORF deletion (Humby and O’Connor, 2015; Krishna et al., 2019) or stop codon insertion (described herein), respectively. Additionally, Krishna et al published pUS28’s requirement for latent monocyte infection using the Titan strain (Krishna et al., 2017). These consistent findings across strains suggest the viral background most likely does not impact pUS28’s requirement for latency. However, conditions used in various latency models may have an impact. A variety of culture systems for the study of latency are characterized (Collins-McMillen et al., 2018; Poole et al., 2019). We use human hematopoietic-derived cells for our experiments (Humby and O’Connor, 2015; Krishna et al., 2019), including the CD34^+^ cell line, Kasumi-3 (O’Connor and Murphy, 2012), THP-1 monocytes (Sinclair et al., 1992), and primary cord blood-derived CD34^+^ hematopoietic cells (Goodrum et al., 2002; Goodrum et al., 2004). Indeed, others have used THP-1 cells (Wu and Miller, 2016; Krishna et al., 2017), as well as primary monocytes (Krishna et al., 2017), to detail pUS28’s ability to repress IE gene and protein expression (Wu and Miller, 2016; Krishna et al., 2017), as well as maintain latency (Krishna et al., 2017). The Crawford et al. study used a slightly different model system: CD34^+^ cells derived from fetal liver (Crawford et al., 2019). It is possible, therefore, that while these cells fully support HCMV latency and reactivation, the underlying biological mechanisms the virus uses are distinct from those it employs in cells of hematopoietic origin. It would prove interesting to determine the outcome of infecting the fetal liver-derived CD34^+^ cells with our viral constructs in the future, which may reveal novel differences, while highlighting similarities, with regards to the function of this key protein in different setting.

In sum, our work presented herein reveals pUS28 expression is critical to HCMV latency. Further, our data reveal the deletion of the *US28* ORF from our constructs does not impact the expression of the polycistronic transcript, *US27*, or that of the downstream *US29* gene. While we and others have begun to interrogate the signaling pathways pUS28 potentiates to maintain viral latency, further work aimed at understanding the cellular and viral factors pUS28 manipulates during this phase of infection will provide insight into the HCMV-host relationship.

## 6 Author Contributions

BK, AW, RS, and CO generated reagents and performed experiments. BK, AW, RS, and CO analyzed the data. CO wrote the manuscript. All authors contributed to manuscript revision and approved the submitted version.

## 8 Acknowledgments

The authors would like to thank Eain A. Murphy, PhD for critical reading of this manuscript and helpful discussions.

## References

Arvin, A.M., Fast, P., Myers, M., Plotkin, S., Rabinovich, R., and National Vaccine Advisory, C. (2004). Vaccine development to prevent cytomegalovirus disease: report from the National Vaccine Advisory Committee. Clin Infect Dis 39(2), 233–239. doi: 10.1086/421999.

Auger, G.A., Pease, J.E., Shen, X., Xanthou, G., and Barker, M.D. (2002). Alanine scanning mutagenesis of CCR3 reveals that the three intracellular loops are essential for functional receptor expression. Eur J Immunol 32(4), 1052–1058. doi: 10.1002/1521-4141(200204)32:4<1052::AID-IMMU1052>3.0.CO;2-L

Balazs, Z., Tombacz, D., Szucs, A., Csabai, Z., Megyeri, K., Petrov, A.N., et al. (2017). Long-Read Sequencing of Human Cytomegalovirus Transcriptome Reveals RNA Isoforms Carrying Distinct Coding Potentials. Sci Rep 7(1), 15989. doi: 10.1038/s41598-017-16262-z.

Bechtel, J.T., and Shenk, T. (2002). Human cytomegalovirus UL47 tegument protein functions after entry and before immediate-early gene expression. J Virol 76(3), 1043–1050. doi: 10.1128/jvi.76.3.1043-1050.2002.

Casarosa, P., Waldhoer, M., LiWang, P.J., Vischer, H.F., Kledal, T., Timmerman, H., et al. (2005). CC and CX3C chemokines differentially interact with the N terminus of the human cytomegalovirus-encoded US28 receptor. J Biol Chem 280(5), 3275–3285. doi: 10.1074/jbc.M407536200.

Cheng, S., Caviness, K., Buehler, J., Smithey, M., Nikolich-Zugich, J., and Goodrum, F. (2017). Transcriptome-wide characterization of human cytomegalovirus in natural infection and experimental latency. Proc Natl Acad Sci U S A 114(49), E10586–E10595. doi: 10.1073/pnas.1710522114.

Collins-McMillen, D., Buehler, J., Peppenelli, M., and Goodrum, F. (2018). Molecular Determinants and the Regulation of Human Cytomegalovirus Latency and Reactivation. Viruses 10(8). doi: 10.3390/v10080444.

Crawford, L.B., Caposio, P., Kreklywich, C., Pham, A.H., Hancock, M.H., Jones, T.A., et al. (2019). Human Cytomegalovirus US28 Ligand Binding Activity Is Required for Latency in CD34(+) Hematopoietic Progenitor Cells and Humanized NSG Mice. MBio 10(4). doi: 10.1128/mBio.01889-19.

Dunn, W., Chou, C., Li, H., Hai, R., Patterson, D., Stolc, V., et al. (2003). Functional profiling of a human cytomegalovirus genome. Proc Natl Acad Sci U S A 100(24), 14223–14228. doi: 10.1073/pnas.2334032100.

Elder, E., and Sinclair, J. (2019). HCMV latency: what regulates the regulators? Med Microbiol Immunol 208(3–4), 431–438. doi: 10.1007/s00430-019-00581-1.

Elder, E.G., Krishna, B.A., Williamson, J., Lim, E.Y., Poole, E., Sedikides, G.X., et al. (2019). Interferon-Responsive Genes Are Targeted during the Establishment of Human Cytomegalovirus Latency. MBio 10(6). doi: 10.1128/mBio.02574-19.

Goodrum, F., Jordan, C.T., Terhune, S.S., High, K., and Shenk, T. (2004). Differential outcomes of human cytomegalovirus infection in primitive hematopoietic cell subpopulations. Blood 104(3), 687–695. doi: 10.1182/blood-2003-12-4344.

Goodrum, F.D., Jordan, C.T., High, K., and Shenk, T. (2002). Human cytomegalovirus gene expression during infection of primary hematopoietic progenitor cells: a model for latency. Proc Natl Acad Sci U S A 99(25), 16255–16260. doi: 10.1073/pnas.252630899.

Griffiths, P., Baraniak, I., and Reeves, M. (2015). The pathogenesis of human cytomegalovirus. J Pathol 235(2), 288–297. doi: 10.1002/path.4437.

Halazonetis, T.D., Georgopoulos, K., Greenberg, M.E., and Leder, P. (1988). c-Jun dimerizes with itself and with c-Fos, forming complexes of different DNA binding affinities. Cell 55(5), 917–924. doi: 10.1016/0092-8674(88)90147-x.

Haskell, C.A., Cleary, M.D., and Charo, I.F. (1999). Molecular uncoupling of fractalkine-mediated cell adhesion and signal transduction. Rapid flow arrest of CX3CR1-expressing cells is independent of G-protein activation. J Biol Chem 274(15), 10053–10058. doi: 10.1074/jbc.274.15.10053.

Humby, M.S., and O’Connor, C.M. (2015). Human Cytomegalovirus US28 Is Important for Latent Infection of Hematopoietic Progenitor Cells. J Virol 90(6), 2959–2970. doi: 10.1128/JVI.02507-15.

Khanna, R., and Diamond, D.J. (2006). Human cytomegalovirus vaccine: time to look for alternative options. Trends Mol Med 12(1), 26–33. doi: 10.1016/j.molmed.2005.11.006.

Krishna, B.A., Humby, M.S., Miller, W.E., and O’Connor, C.M. (2019). Human cytomegalovirus G protein-coupled receptor US28 promotes latency by attenuating c-fos. Proc Natl Acad Sci U S A 116(5), 1755–1764. doi: 10.1073/pnas.1816933116.

Krishna, B.A., Miller, W.E., and O’Connor, C.M. (2018). US28: HCMV’s Swiss Army Knife. Viruses 10(8). doi: 10.3390/v10080445.

Krishna, B.A., Poole, E.L., Jackson, S.E., Smit, M.J., Wills, M.R., and Sinclair, J.H. (2017). Latency-Associated Expression of Human Cytomegalovirus US28 Attenuates Cell Signaling Pathways To Maintain Latent Infection. MBio 8(6). doi: 10.1128/mBio.01754-17.

Ljungman, P., Boeckh, M., Hirsch, H.H., Josephson, F., Lundgren, J., Nichols, G., et al. (2017). Definitions of Cytomegalovirus Infection and Disease in Transplant Patients for Use in Clinical Trials. Clin Infect Dis 64(1), 87–91. doi: 10.1093/cid/ciw668.

Maussang, D., Langemeijer, E., Fitzsimons, C.P., Stigter-van Walsum, M., Dijkman, R., Borg, M.K., et al. (2009). The human cytomegalovirus-encoded chemokine receptor US28 promotes angiogenesis and tumor formation via cyclooxygenase-2. Cancer Res 69(7), 2861–2869. doi: 10.1158/0008-5472.CAN-08-24871.

Maussang, D., Verzijl, D., van Walsum, M., Leurs, R., Holl, J., Pleskoff, O., et al. (2006). Human cytomegalovirus-encoded chemokine receptor US28 promotes tumorigenesis. Proc Natl Acad Sci U S A 103(35), 13068–13073. doi: 10.1073/pnas.0604433103.

Miller, W.E., Zagorski, W.A., Brenneman, J.D., Avery, D., Miller, J.L., and O’Connor, C.M. (2012). US28 is a potent activator of phospholipase C during HCMV infection of clinically relevant target cells. PLoS One 7(11), e50524. doi: 10.1371/journal.pone.0050524.

Murphy, E., Vanicek, J., Robins, H., Shenk, T., and Levine, A.J. (2008). Suppression of immediate-early viral gene expression by herpesvirus-coded microRNAs: implications for latency. Proc Natl Acad Sci U S A 105(14), 5453–5458. doi: 10.1073/pnas.0711910105.

Noriega, V.M., Gardner, T.J., Redmann, V., Bongers, G., Lira, S.A., and Tortorella, D. (2014). Human cytomegalovirus US28 facilitates cell-to-cell viral dissemination. Viruses 6(3), 1202–1218. doi: 10.3390/v6031202.

O’Connor, C.M., and Miller, W.E. (2014). Methods for studying the function of cytomegalovirus GPCRs. Methods Mol Biol 1119, 133–164. doi: 10.1007/978-1-62703-788-4_10.

O’Connor, C.M., and Murphy, E.A. (2012). A myeloid progenitor cell line capable of supporting human cytomegalovirus latency and reactivation, resulting in infectious progeny. J Virol 86(18), 9854–9865. doi: 10.1128/JVI.01278-12.

O’Connor, C.M., and Shenk, T. (2011). Human cytomegalovirus pUS27 G protein-coupled receptor homologue is required for efficient spread by the extracellular route but not for direct cell-to-cell spread. J Virol 85(8), 3700–3707. doi: 10.1128/JVI.02442-10.

O’Connor, C.M., and Shenk, T. (2012). Human cytomegalovirus pUL78 G protein-coupled receptor homologue is required for timely cell entry in epithelial cells but not fibroblasts. J Virol 86(21), 11425–11433. doi: 10.1128/JVI.05900-11.

Poole, E., Huang, C.J.Z., Forbester, J., Shnayder, M., Nachshon, A., Kweider, B., et al. (2019). An iPSC-Derived Myeloid Lineage Model of Herpes Virus Latency and Reactivation. Front Microbiol 10, 2233. doi: 10.3389/fmicb.2019.02233.

Ramanan, P., and Razonable, R.R. (2013). Cytomegalovirus infections in solid organ transplantation: a review. Infect Chemother 45(3), 260–271. doi: 10.3947/ic.2013.45.3.260.

Rovati, G.E., Capra, V., and Neubig, R.R. (2007). The highly conserved DRY motif of class A G protein-coupled receptors: beyond the ground state. Mol Pharmacol 71(4), 959–964. doi: 10.1124/mol.106.029470.

Schwartz, T.W., Frimurer, T.M., Holst, B., Rosenkilde, M.M., and Elling, C.E. (2006). Molecular mechanism of 7TM receptor activation--a global toggle switch model. Annu Rev Pharmacol Toxicol 46, 481–519. doi: 10.1146/annurev.pharmtox.46.120604.141218.

Shnayder, M., Nachshon, A., Krishna, B., Poole, E., Boshkov, A., Binyamin, A., et al. (2018). Defining the Transcriptional Landscape during Cytomegalovirus Latency with Single-Cell RNA Sequencing. MBio 9(2). doi: 10.1128/mBio.00013-18.

Silva, M.C., Yu, Q.C., Enquist, L., and Shenk, T. (2003). Human cytomegalovirus UL99-encoded pp28 is required for the cytoplasmic envelopment of tegument-associated capsids. J Virol 77(19), 10594–10605. doi: 10.1128/jvi.77.19.10594-10605.2003.

Sinclair, J.H., Baillie, J., Bryant, L.A., Taylor-Wiedeman, J.A., and Sissons, J.G. (1992). Repression of human cytomegalovirus major immediate early gene expression in a monocytic cell line. J Gen Virol 73 (Pt 2), 433–435. doi: 10.1099/0022-1317-73-2-433.

Sinzger, C., Hahn, G., Digel, M., Katona, R., Sampaio, K.L., Messerle, M., et al. (2008). Cloning and sequencing of a highly productive, endotheliotropic virus strain derived from human cytomegalovirus TB40/E. J Gen Virol 89(Pt 2), 359–368. doi: 10.1099/vir.0.83286-0.

Slinger, E., Maussang, D., Schreiber, A., Siderius, M., Rahbar, A., Fraile-Ramos, A., et al. (2010). HCMV-encoded chemokine receptor US28 mediates proliferative signaling through the IL-6-STAT3 axis. Sci Signal 3(133), ra58. doi: 10.1126/scisignal.2001180.

Umashankar, M., and Goodrum, F. (2014). Hematopoietic long-term culture (hLTC) for human cytomegalovirus latency and reactivation. Methods Mol Biol 1119, 99–112. doi: 10.1007/978-1-62703-788-4_7.

Waldhoer, M., Kledal, T.N., Farrell, H., and Schwartz, T.W. (2002). Murine cytomegalovirus (CMV) M33 and human CMV US28 receptors exhibit similar constitutive signaling activities. J Virol 76(16), 8161–8168. doi: 10.1128/jvi.76.16.8161-8168.2002.

Wu, S.E., and Miller, W.E. (2016). The HCMV US28 vGPCR induces potent Galphaq/PLC-beta signaling in monocytes leading to increased adhesion to endothelial cells. Virology 497, 233–243. doi: 10.1016/j.virol.2016.07.025.

Yu, D., Silva, M.C., and Shenk, T. (2003). Functional map of human cytomegalovirus AD169 defined by global mutational analysis. Proc Natl Acad Sci U S A 100(21), 12396–12401. doi: 10.1073/pnas.1635160100.

Zhu, H., Shen, Y., and Shenk, T. (1995). Human cytomegalovirus IE1 and IE2 proteins block apoptosis. J Virol 69(12), 7960–7970.

