## Supplemental Data for "The Requirement For US28 During Cytomegalovirus Latency Is Independent Of US27 And US29 Gene Expression"

**Supplementary Data**

Supplementary Tables: 1

Supplementary Figures: 3

**Supplementary Table 1.** Oligonucleotides used in this study.

| Primer Use | Primer Direction | Primer Sequence |
|---|---|---|
| **3xF-Kan-frt insertion** | forward§ | TCTTCCGACACGCTGTCCGACGAGGTGTGTCGCGTCTCACAAATTATA CCGTTAGATTATAAAGATGATGATGATAAA |
| | reverse§ | AGAGGGGCGGACACGGGGTTTGTATGAAAAGGCCGAGGTAGCGCTTT TTTAGGCCGCGGGAATTCGAAGTT |
| ***galK* insertion** | forward¶ | GCACCGAGGGCAGAACTGGTGCTATCATGACACCGACGACGACGACC GCGcctgttgacaattaatcatcggca |
| | reverse¶ | TGAAAACACAAGGAGTCGCGTCTTCATCGTAGTCAAACTCCGTCGTGA GTTtcagcactgtcctgctcctt |
| ***galK* reversion oligo** | top | CACCGAGGGCAGAACTGGTGCTATCATGACACCGACGACGACGACCG CGTAACTCACGACGGAGTTTGACTACGATGAAGACGCGACTCCTTGTG TTTT |
| | bottom | AAAACACAAGGAGTCGCGTCTTCATCGTAGTCAAACTCCGTCGTGAGT TACGCGGTCGTCGTCGTCGGTGTCATGATAGCACCAGTTCTGCCCTCG GTG |
| **Multi-tag vGPCR virus – *galK* insertion** | | |
| UL33-c-myc | forward¶ | ACAAAAATCCCCCATCGACTCTCACAATCGCATCATAACCTCAGCGGG GTAcctgttgacaattaatcatcggca |
| | reverse¶ | AAATGGCGACGGGTTCTGGTGCTTTCTGAATAAAGTAACAGGAAAGCT CAtcagcactgtcctgctcctt |
| UL78-V5 | forward¶ | TGCACCGACGGCGAAAACACCGTCGCGTCCGACGCAACGGTGACGGC ATTAcctgttgacaattaatcatcggca |
| | reverse¶ | GTGATTTATCTGCCACTTTTCTCCCCGCTGCCGTACAGCGCCGCCGCTC Atcagcactgtcctgctcctt |
| US27-3xHA | forward¶ | TATGACAGAAAAAATGCACCTATGGAGTCCGGGGAGGAGGAATTTCT ATTGcctgttgacaattaatcatcggca |
| | reverse¶ | CAATGAGCAAAAATAGATGTGCGGCGGACGCGTGAAAGAGGATCGAA |

| | | |
|---|---|---|
| | | TTAtcagcactgtcctgctcctt |
| **Multi-tag vGPCR virus – *galK* reversion gBlocks** | | |
| UL33-c-myc | | ACAAAAATCCCCCATCGACTCTCACAATCGCATCATAACCTCAGCGGG GTAGAACAAAAACTTATTTCTGAAGAAGATCTTTGAGCTTTCCTGTTAC TTTATTCAGAAAGCACCAGAACCCGTCGCCATTT |
| UL78-V5 | | TGCACCGACGGCGAAAACACCGTCGCGTCCGACGCAACGGTGACGGC ATTAGGTAAGCCTATCCCTAACCCTCTCCTCGGTCTCGATTCTACGTGA GCGGCGGCGCTGTACGGCAGCGGGGAGAAAAGTGGCAGATAAATCAC |
| US27-3xHA | | TATGACAGAAAAAATGCACCTATGGAGTCCGGGGAGGAGGAATTTCT ATTGTACCCATATGACGTTCCAGACTACGCGTATCCGTACGACGTTCC GGATTACGCTTACCCTTACGACGTACCTGACTACGCTTAATTCGATCCT CTTTCACGCGTCCGCCGCACATCTATTTTTGCTCATTG |
| **Multi-tag vGPCR virus – *galK* reversion gBlock amplifying primers** | | |
| UL33-c-myc | forward | ACAAAAATCCCCCATCGACTC |
| | reverse | AAATGGCGACGGGTTCTGGTG |
| UL78-V5 | forward | TGCACCGACGGCGAAAACACC |
| | reverse | GTGATTTATCTGCCACTTTTC |
| US27-3xHA | forward | TATGACAGAAAAAATGCACCT |
| | reverse | CAATGAGCAAAAATAGATGTG |
| **RTqPCR** | | |
| US27 | forward | CCGTATGGTGCGGTTTATCATTA |
| | reverse | CTAAAAATAGCGCCAGGTTGAAAGG |
| US28 | forward | CCAGAATCGTTGCGGTGTCTCAGT |
| | reverse | CGTGTCCACAAACAGCGTCAGGT |
| US29 | forward | CGACGAGACAACAATGAC |

| | | |
|---|---|---|
| | reverse | AATTGACGGTCCACTGAG |
| UL123 | forward | GCCTTCCCTAAGACCACCAAT |
| | reverse | ATTTTCTGGGCATAAGCCATAATC |
| UL99 | forward | GTGTCCCATTCCCGACTCG |
| | reverse | TTCACAACGTCCACCCACC |
| GAPDH | forward | ACCCACTCCTCCACCTTTGAC |
| | reverse | CTGTTGCTGTAGCCAAATTCGT |

§ Underlined sequences are complementary to the pKan-frt template.
¶ Lowercase sequences are complementary to the pGalK template.

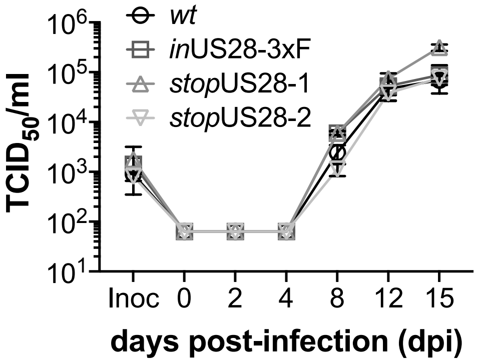


**Supplementary Figure 1. US28 recombinant viruses display wild type growth kinetics in lytically-infected primary fibroblasts.** MRC-5 cells were infected as indicated (moi = 0.01). Supernatants from each culture were collected at the indicated days post-infection (dpi). Infectious virus was quantified by $TCID_{50}$ on naïve fibroblasts. Inoc, inoculum. All samples were analyzed in triplicate. Error bars indicate standard deviation, and statistical significance was calculated using two-way ANOVA analyses followed by Tukey's post-hoc analyses. Values were not statistically significant.

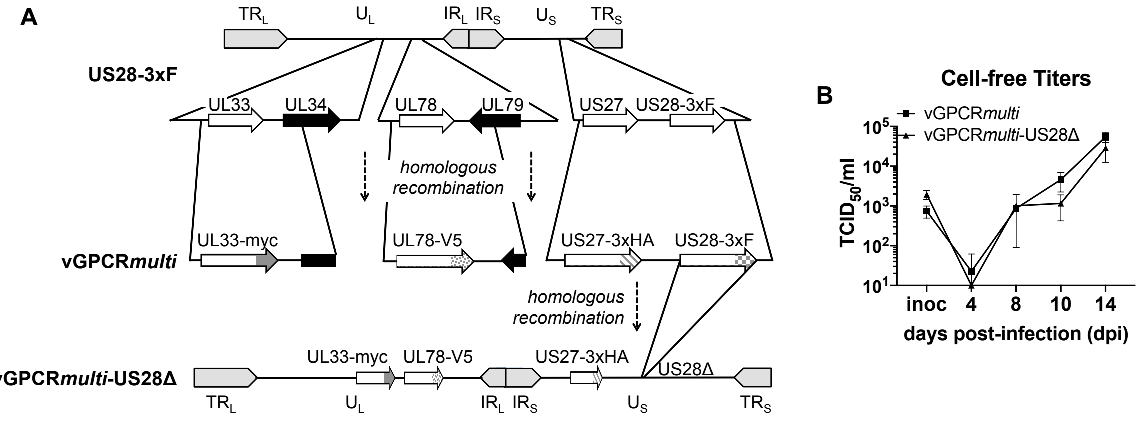


**Supplementary Figure 2. TB40/E*mCherry*-vGPCR*multi* and- vGPCR*multi*-US28Δ replicate to wild type titers in lytically infected fibroblasts. (A)** TB40/E*mCherry*-US28-3xF (Miller et al., 2012) was used to generate TB40/E*mCherry*-vGPCR*multi* (vGPCR*multi*), containing epitope tags on each additional vGPCR, as depicted. vGPCR*multi* was subsequently used to generate vGPCR*multi*-US28Δ, which lacks the entire US28 ORF, inclusive of the triple FLAG epitope tag. **(B)** Multi-step growth analyses of the viral recombinants in **(A)** was evaluated in lytically infected NuFF-1 fibroblasts (moi = 0.01). At the indicated times post-infection, infectious supernatants were collected, and titers were determined using $TCID_{50}$ assay on naïve NuFF-1 fibroblasts in triplicate. Inoc, inoculum. Error bars indicate standard deviation, and statistical significance was calculated using two-way ANOVA with Sidak's post-hoc analysis. Values were not statistically significant.

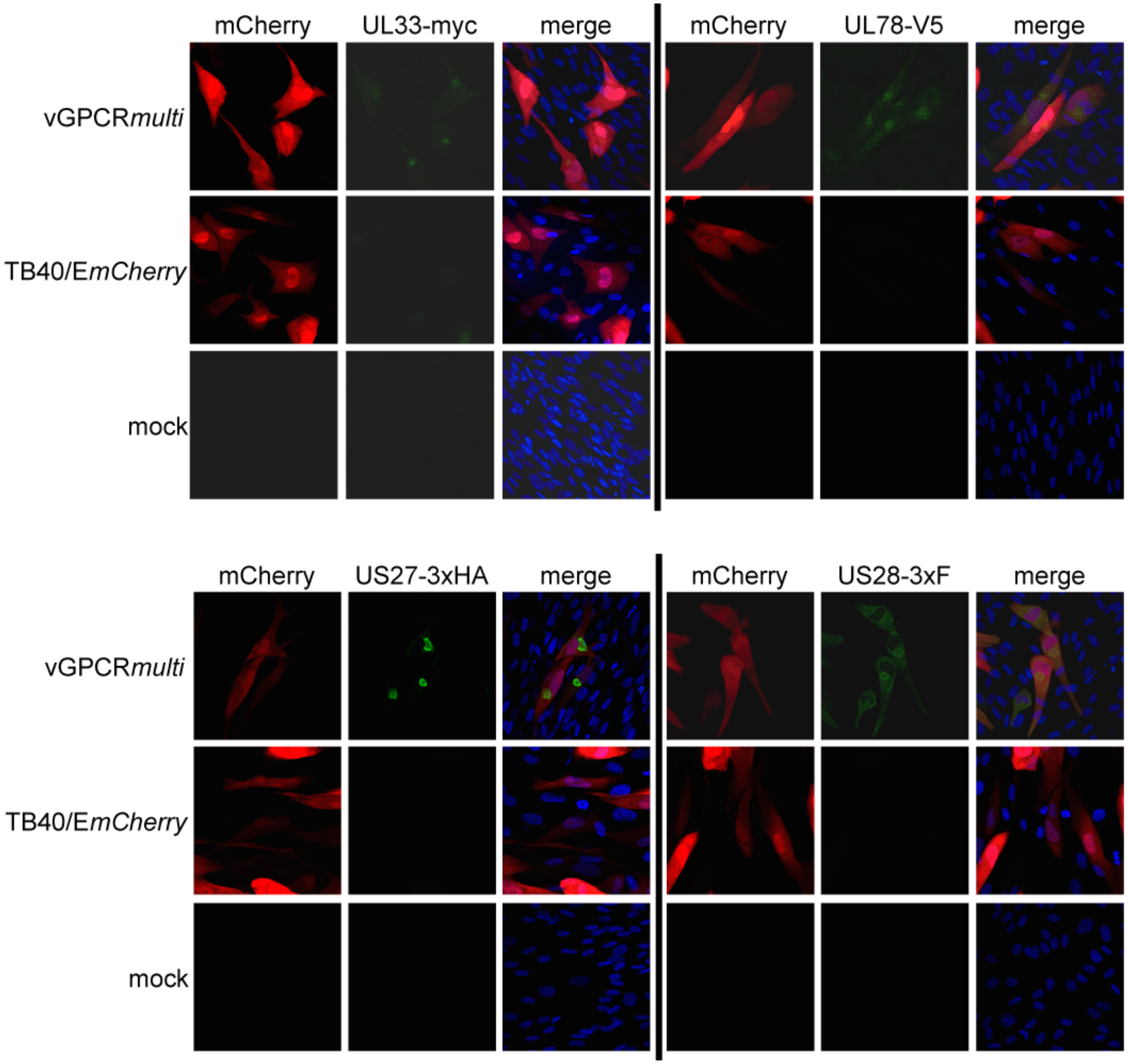


**Supplementary Figure 3. TB40/E*mCherry*-vGPCR*multi* expresses each vGPCR following lytic infection.** NuFF-1 fibroblasts were mock-infected or lytically infected (moi = 0.5) as shown and harvested 96 hpi for IFA. The appropriate antibodies were used to detect the indicated epitope tag (see Materials and Methods for specific antibody details): UL33 (anti-myc), UL78 (anti-V5), US27 (anti-HA), US28 (anti-FLAG), each shown in green. mCherry (red) is a marker of infection, and nuclei were visualized with DAPI (blue). Images were acquired with a 40x objective with 1.5x magnification. Representative images are shown (n=3).
